# Transcriptional Plasticity of *Trichoderma atroviride* in Response to Different Plant Pathogens

**DOI:** 10.64898/2026.09.15.751736

**Authors:** Etienne Brémand, Franck Bastide, Justine Colou, Nicolas Denancé, Thomas Guillemette

## Abstract

Fungal mycoparasitism is commonly viewed as a broadly conserved lifestyle relying on the secretion of cell wall-degrading enzymes and specialized metabolites. However, whether *Trichoderma* deploys a universal antagonistic program or dynamically adapts its molecular arsenal according to prey identity remains largely unknown. Here, we investigated the phenotypic and transcriptomic responses of four highly antagonistic *Trichoderma atroviride* strains confronted with damping off pathogens from three phylogenetically distinct lineages: the ascomycete *Alternaria brassicicola*, the basidiomycete *Rhizoctonia solani*, and the oomycete *Globisporangium ultimum*. *In vitro* confrontation assays were performed between the four *T. atroviride* strains and nine pathogen strains (three strains per species). Despite notable intraspecific variation in pathogen sensitivity, all four *T. atroviride* strains remained highly effective against all tested pathogens.

Comparative RNA-seq analyses during direct confrontation with one representative strain per pathogen species revealed strong prey-dependent transcriptional plasticity. Interactions with the two fungal pathogens elicited a robust induction of classical mycoparasitic machinery, including glycoside hydrolases, secreted peptidases, and effector-like proteins, whereas these responses were markedly attenuated in interaction with *G. ultimum*. Specialized metabolite biosynthetic genes were broadly induced across all interactions, but with largely distinct gene sets depending on the pathogen, with the strongest divergence observed for confrontation with *G. ultimum*. Beyond this shared response, each fungal pathogen also triggered a distinct host-specific program. In response to *A. brassicicola* and *R. solani*, *T. atroviride* induced distinct sets of genes associated with cell wall organization, while the response to *R. solani* specifically involved the activation of peroxisomal functions and aromatic amino acid biosynthesis pathways. Receptors potentially involved in pathogen perception also showed pathogen-dependent expression patterns, with distinct GPCRs (G protein-coupled receptors) overexpressed depending on the pathogen encountered. NLRs (NOD-like receptors) displayed a similar pattern, although overexpression was detected only in response to *A. brassicicola* and *G. ultimum*.

Together, these results demonstrate that *T. atroviride* does not rely on a fixed mycoparasitic program but instead dynamically remodels its transcriptome according to prey identity, revealing a high degree of transcriptional plasticity underlying broad-spectrum antagonism.

## 1 Introduction

The growing awareness of the environmental and health impacts associated with chemical pesticides has accelerated the search for sustainable alternatives capable of preserving biodiversity, soil quality, and human health. Among these alternatives, biological control strategies based on the use of beneficial microorganisms have emerged as promising tools for plant protection. Within this context, fungi belonging to the genus *Trichoderma* are among the most widely studied and commercially exploited biocontrol agents due to their remarkable ability to antagonize, parasitize, and consume a broad range of plant pathogens (Dutta et al., 2022).

Among them, *Trichoderma atroviride* has demonstrated strong biocontrol activity both *in vitro* and *in planta* against numerous phytopathogens, including ascomycete fungi such as *Alternaria*, *Botrytis* and *Fusarium* (Brémand et al., 2026b; Chateau et al., 2024; Pascouau et al., 2023); basidiomycete fungi such as *Rhizoctonia* and *Armillaria* (Atanasova et al., 2013; Chen et al., 2023); as well as oomycetes including *Phytophthora*, *Pythium* and *Globisporangium* (Reithner et al., 2011; Chen et al., 2024). In addition, *T. atroviride* can also target other plant-associated pests, including bacteria (Papaianni et al., 2020), nematodes (Pinto et al., 2026; Sharon et al., 2007) and insects (Contreras-Cornejo et al., 2018; Poveda, 2021), highlighting its exceptionally broad host spectrum.

The molecular mechanisms underlying mycoparasitism in *T. atroviride* have been extensively investigated through genomic and transcriptomic approaches. These mechanisms primarily involve the secretion of cell wall-degrading enzymes (CWDEs), such as chitinases, glucanases, and peptidases, which enable the degradation of prey cell walls, together with the production of specialized metabolites that inhibit pathogen growth (Brémand et al., 2026a). However, increasing evidence suggests that these antagonistic mechanisms are not universally conserved. Previous studies have shown substantial variation both between *Trichoderma* species (Atanasova et al., 2013) and between strains of *T. atroviride* (Brémand et al., 2026a). Yet, despite these observations, most transcriptomic studies have focused on single interactions involving one *Trichoderma* strain confronted with one pathogen species (Chen et al., 2023; Steindorff et al., 2014; Wang et al., 2024, 2023a, 2023b), leaving unresolved whether *Trichoderma* deploys a conserved mycoparasitic program or dynamically adjusts its molecular arsenal depending on prey identity.

Our previous study further highlighted that the antagonistic efficiency of *Trichoderma* strongly depends on both phylogenetic background and pathogen identity (Brémand et al., 2026b). Strains belonging to the *Longibrachiatum* clade were generally more efficient against the oomycete *Globisporangium ultimum*, whereas strains from the *Viride* clade displayed higher antagonistic activity against the fungal pathogens *Alternaria brassicicola* and *Rhizoctonia solani*. Phylogenetically distinct *Trichoderma* lineages may rely on different molecular strategies, which could influence their efficiency against different pathogens and contribute to their distinct host ranges (Atanasova et al., 2013).

In the present study, we focused on four *T. atroviride* strains exhibiting strong antagonistic activity against three damping off pathogens (*A. brassicicola*, *R. solani*, and *G. ultimum*). By combining phenotypic characterization and comparative transcriptomics during direct confrontation, we aimed to identify the molecular mechanisms underlying this broad antagonistic spectrum. More specifically, we investigated whether these strains rely on a conserved set of broadly effective mechanisms or whether transcriptional plasticity enables the dynamic remodeling of their antagonistic arsenal according to prey identity.

## 2 Materials and Methods

### 2.1 Strains and Cultivation Conditions

*Trichoderma atroviride* strains were isolated from flax seeds (P3080) or lettuce seeds (P3116) by GEVES (Angers, France), or from marine environments (MMS1295 and N1508) by ISOMer (Nantes, France). Nine damping-off pathogens were used in this study. *Alternaria brassicicola* strains Abra43 (CIRM-CF, BRFM3693), BA 1370, and BA 1374 were isolated from radish leaves, cabbage leaves, and cabbage seeds, respectively, by IRHS (Angers, France). *Rhizoctonia solani* strains PAT-002, PAT-006, and PAT-009 were isolated from lettuce, corn salad, and radish, respectively, by CDDM (Nantes, France). *Globisporangium ultimum* var. *ultimum* strains CBS 114.19, CBS 378.34, and CBS 725.94, isolated from gymnosperm seedlings, *Trifolium pratense*, and wheat, respectively, were obtained from the Westerdijk Fungal Biodiversity Institute (Utrecht, the Netherlands). For long-term storage, all strains were maintained at −80°C as mycelial explants stored in 30% glycerol, except *G. ultimum*, which was preserved at 4°C as mycelial explants stored in sterile water. For fresh cultures, all strains were grown on Potato Dextrose Agar (PDA, Biokar, 39 g/L) at 22°C in the dark.

### 2.2 *In Vitro* Confrontation Assays

A dual culture assay was performed in square Petri dishes (120 mm × 120 mm) containing PDA medium. Plates were inoculated with two 5-mm-diameter mycelial plugs placed at opposite corners of the dish (one from the pathogen and one from *T. atroviride*). Plugs of *A. brassicicola* were placed 3 days before *T. atroviride*, whereas plugs of *R. solani* and *G. ultimum* were inoculated simultaneously with *T. atroviride* due to differences in growth rates. Three replicates were performed for each condition, and the assay was repeated in three independent experiments. Plates were incubated at 20°C in the dark. Ten days after pathogen contact, the mycoparasitic performance of each *T. atroviride* strain was visually assessed using the scale from Brémand et al. (2026b). Non-parametric Kruskal-Wallis tests were applied to evaluate global significant differences (p < 0.05) for *Trichoderma* strains and pathogen isolates. Multiple pairwise comparisons were conducted using Dunn’s post hoc test, and p-values were adjusted using the Benjamini-Hochberg (BH) false discovery rate method. Distinct letters indicate significant differences (p < 0.05) according to the compact letter display.

### 2.3 Measurement of Pathogen Growth Inhibition by *T. atroviride* Culture Filtrates

Liquid cultures of *T. atroviride* and their filtration were performed according to Chateau et al. (2024). For *A. brassicicola*, 10^6^ conidia·mL^-1^ were collected from a 7-day-old culture on V8 agar medium and suspended in sterile water. For *R. solani*, a mycelial suspension was used as inoculum as described by Brémand et al. (2026a) and the suspension was adjusted to 10⁴ CFU·mL^-1^. For *G. ultimum*, 10^4^ oospores·mL^-1^ were harvested from 15-days old cultures on Synthetischer Nährstoffarmer Agar (SNA) medium (Nirenberg, 1976). The sensitivity of the pathogens to *T. atroviride* culture filtrates was tested in liquid medium using 96-well plates and a 635-nm laser nephelometer (NEPHELOstar, BMG Labtech) (Joubert et al., 2010). PDB (Potato Dextrose Broth, Difco, 24 g·L^-1^) was inoculated with 10% (v/v) of pathogen suspension and 10% (v/v) of *T. atroviride* culture filtrate in a final volume of 200 μL. Plates were incubated at 25°C and growth was monitored every 10 minutes for 35 hours for *A. brassicicola* and *G. ultimum*, and 96 hours for *R. solani*. Three replicates were performed for each condition, and the assay was repeated in three independent experiments. The area under the curve (AUC) was calculated for each growth curve. Non-parametric Kruskal-Wallis tests were applied to evaluate global significant differences (p < 0.05) for *Trichoderma* strains and pathogen isolates. Multiple pairwise comparisons were conducted using Dunn’s post hoc test, and p-values were adjusted using the Benjamini-Hochberg (BH) false discovery rate method. Distinct letters indicate significant differences (p < 0.05) according to the compact letter display. The percentage of inhibition was determined for each condition containing a *T. atroviride* filtrate relative to the control condition without filtrate.

### 2.4 RNA Sequencing and Differential Expression Analyses

*T. atroviride* transcriptomes were analyzed during *in vitro* confrontation assays using the same experimental design as described in section 2.2, but adapted with PDA medium covered with a pectocellulosic membrane. To ensure synchronized contact, the initial spacing between *T. atroviride* and each pathogen was adjusted according to their growth rates and plugs of *A. brassicicola* Abra43, *R. solani* PAT-009, and *G. ultimum* CBS 114.19 were placed 7 days before, 1 day after, or 2 days after *T. atroviride*, respectively. Self-confrontation of *T. atroviride* (second plug added one day after the first) served as a control. All assays were performed in triplicate. Samples were collected from the colony edge of *T. atroviride* at two time points: before physical contact (four days after *T. atroviride* inoculation) and after contact (seven days after *T. atroviride* inoculation). RNA extraction and sequencing were performed as described by Brémand et al. (2026a).

Reads were aligned to the *T. atroviride* N1508 transcriptome (GenBank accession number GCA_982374845.1) using Salmon v1.10.3 (Patro et al., 2017) to obtain counts and TPM (Transcripts Per Million) values. A differential expression analysis was performed using the R package AskoR v1.0.0 (Carvalho et al., 2021). This package relies on edgeR v4.0.3 (Chen et al., 2025) and defines genes as significantly differentially expressed when the absolute log2 fold change is > 1 and the FDR (False Discovery Rate) < 0.05. Comparisons were performed between pre- and post-contact conditions using combined datasets from the four *T. atroviride* strains for each pathogen, as well as for the self-confrontation condition.

Different UpSet plots were constructed using the ggupset v0.4.1 and ggplot2 v4.0.0 packages to identify genes differentially expressed in *T. atroviride* that are shared across responses to the different pathogens. Additional UpSet analyses were performed on specific gene categories. Specialized metabolite biosynthetic gene clusters were predicted using fungiSMASH v8.0.4 (Blin et al., 2025). Glycoside hydrolases were identified through CAZyme annotation performed using dbCAN3 v12 (Zheng et al., 2023). Peptidases were annotated by performing BLASTp searches of the predicted proteome against the MEROPS peptidase database v12.4 (Rawlings et al., 2010). SignalP v5.0 (Almagro Armenteros et al., 2019) was used to predict secreted proteins, enabling the identification of secreted peptidases as well as SSCPs (Small Secreted Cysteine-rich Proteins). SSCPs were defined as predicted secreted proteins shorter than 300 amino acids and containing more than three cysteine residues. NLR (NOD-like receptors) were identified as described by Dyrka et al. (2014) through the detection of protein domains using the PFAM (Protein families database) domain (Mistry et al., 2021). PFAM annotations were obtained using eggNOG-mapper v2.1.13 (Cantalapiedra et al., 2021). Gene Ontology terms were retrieved through the STRING database v12.0 (Szklarczyk et al., 2025).

## 3 Results

### 3.1 Mycoparasitic Performance of *T. atroviride* Against Different Plant Pathogens

*In vitro* confrontation assays showed a strong parasitic ability for all four *T. atroviride* strains. P3080 and P3116 exhibited parasitism intensities of 87% and 88%, respectively, compared to 95% and 94% for MMS1295 and N1508. When comparing results across pathogens, no significant differences were observed among the three selected strains of *A. brassicicola*. For *R. solani*, strain PAT-006 was significantly more susceptible to *T. atroviride* than the two other strains. Among the *G. ultimum* strains, CBS 378.34 was significantly more resistant to *T. atroviride* than the other strains. Overall, parasitism levels were high across all strains, generally exceeding 90%, except for *R. solani* PAT-002 and PAT-009, which showed lower parasitism levels of 78% and 76%, respectively.

Antibiosis assays using nephelometry were performed to determine the growth inhibition percentage of the nine pathogens in response to culture filtrates from the four *T. atroviride* strains. All four strains produced highly effective culture filtrates, with inhibition rates exceeding 80%. At the pathogen level, no significant differences were observed among the three *A. brassicicola* strains tested. For *R. solani*, strain PAT-006, the most susceptible in the *in vitro* confrontation assays, was unexpectedly the most resistant to *T. atroviride* culture filtrates. For *G. ultimum*, although strain CBS 378.34 showed higher resistance in confrontation assays, no differences in resistance to *T. atroviride* culture filtrates were observed compared with the other strains.

**Figure 1:**
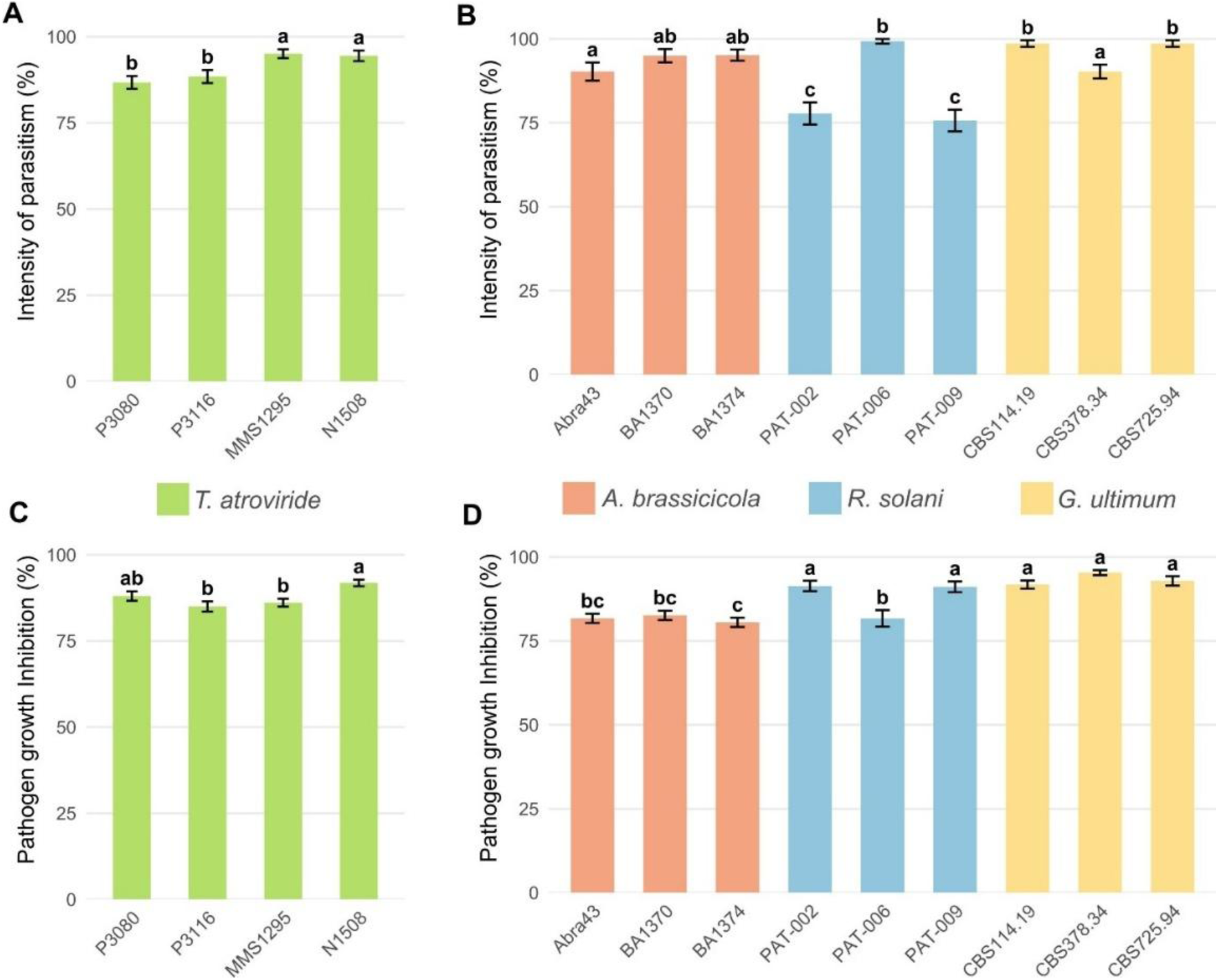
Antagonistic activity of four *Trichoderma atroviride* strains against nine phytopathogenic agents. Top panels show the results of *in vitro* confrontation assays, with parasitism intensity corresponding to the scoring scale (A and B). Bottom panels show antibiosis assays, with percentages of pathogen growth inhibition according to the culture filtrates of *T. atroviride* (C and D). On the left, results for the nine pathogens were averaged for each *T. atroviride* strain (A and C), whereas on the right, results for the four *T. atroviride* strains were averaged for each pathogen (B and D). Error bars represent the standard error. Different letters indicate significant differences between conditions according to a Kruskal-Wallis test followed by Dunn’s post-hoc test (p < 0.05).

Overall, these results indicate that despite measurable intraspecific variation among pathogen strains, *T. atroviride* strains and their respective culture filtrates remain highly effective against all tested pathogens. However, it remains unclear whether the transcriptional responses to these three pathogens differ. To address this question, we performed a transcriptomic analysis of the interactions between the four *T. atroviride* strains and representative strains of each pathogen species.

### 3.2 Pathogen-Dependent Transcriptional Plasticity in *T. atroviride*

The transcriptional responses of the four *T. atroviride* strains were assessed by RNA-seq during *in vitro* confrontation assays against *A. brassicicola* Abra43, *R. solani* PAT-009, and *G. ultimum* CBS 114.19. Differential expression analyses were performed by pooling the transcriptomic data from the four *T. atroviride* strains and comparing gene expression profiles before and after contact with each pathogen, as well as during the control condition consisting of self-confrontation between *T. atroviride* strains.

Contact with *A. brassicicola* and *R. solani* induced the strongest transcriptional reprogramming, with 1,814 and 1,522 upregulated genes, and 627 and 780 downregulated genes, respectively. In contrast, contact with *G. ultimum* resulted in the upregulation of 1,081 genes and the downregulation of 420 genes. Self-confrontation triggered the upregulation of 1,163 genes and the downregulation of 343 genes. Figure 2 illustrates the number of differentially expressed genes detected between before and after contact with each pathogen. Surprisingly, only 88 genes were commonly upregulated in response to all three pathogens, indicating that *T. atroviride* deploys largely distinct transcriptional responses depending on the interacting microorganism.

**Figure 2:**
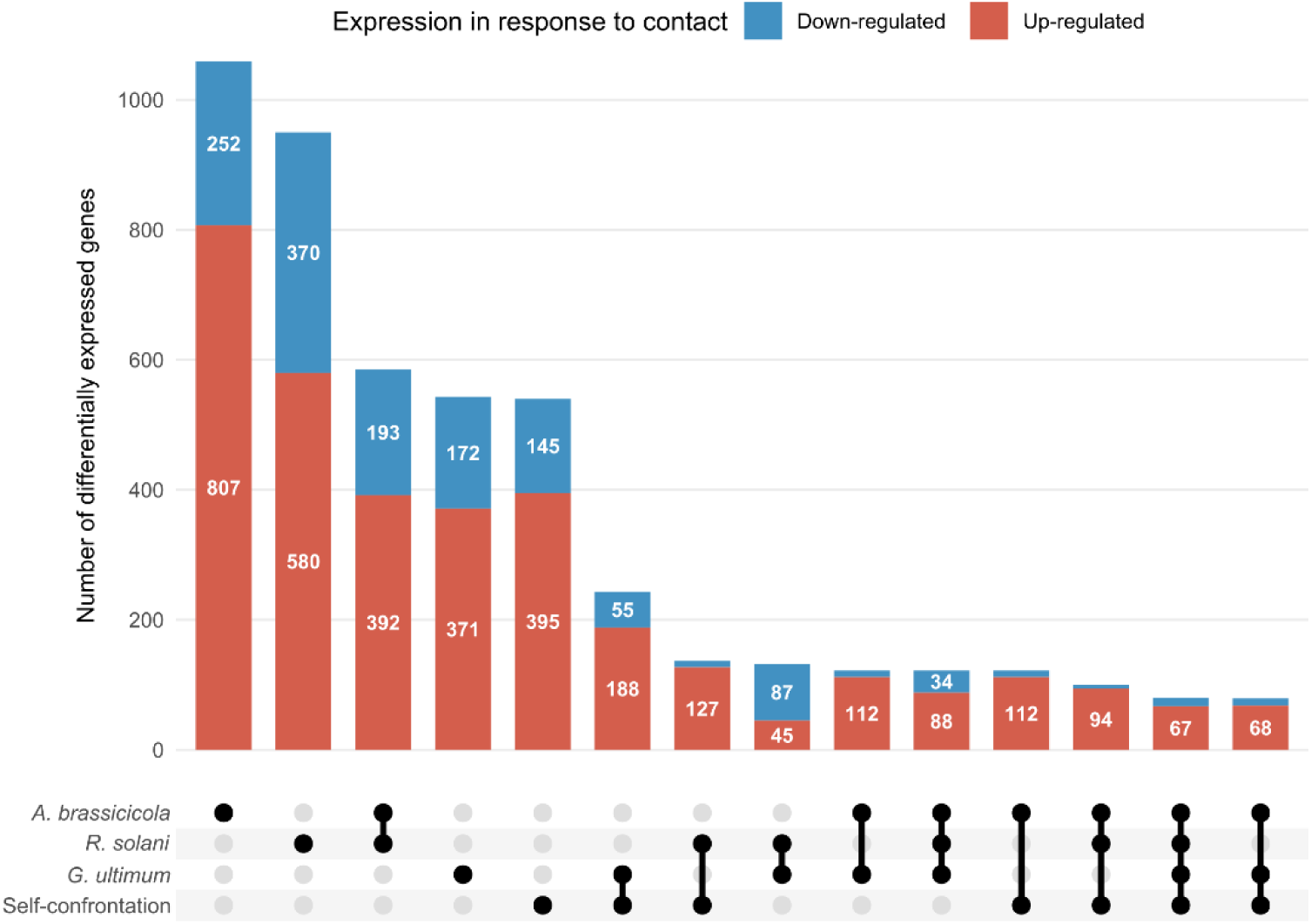
UpSet plot of differentially expressed genes in *Trichoderma atroviride* in response to contact. Genes differentially expressed between before and after contact with *Alternaria brassicicola*, *Rhizoctonia solani*, *Globisporangium ultimum*, or during self-confrontation between *T. atroviride* strains. Genes upregulated in response to contact are shown in red, whereas downregulated genes are shown in blue. Conditions with fewer than 50 differentially expressed genes are not shown.

The responses to the two fungal pathogens, *A. brassicicola* and *R. solani*, were markedly more similar to each other than to the response elicited by the oomycete *G. ultimum*. Indeed, 392 genes were commonly upregulated in response to both *A. brassicicola* and *R. solani* (Figure 2). A substantial proportion of the upregulated genes were pathogen-specific. Specifically, 807 genes were exclusively induced in response to *A. brassicicola* (45% of all upregulated genes in this condition), 580 genes were specific to *R. solani* (38%), and 371 genes were uniquely induced in response to *G. ultimum* (34%). Downregulated genes were less numerous than upregulated genes but exhibited similar pathogen-dependent patterns. Only 34 genes were commonly downregulated in response to all three pathogens, whereas 193 genes were shared between the responses to *A. brassicicola* and *R. solani*. In addition, a large fraction of downregulated genes remained pathogen-specific, with 370, 252, and 172 genes uniquely repressed in response to *A. brassicicola*, *R. solani*, and *G. ultimum*, respectively. Most genes upregulated during *T. atroviride* self-confrontation were also specific to this condition (395 genes). However, overlaps with pathogen-induced responses were also observed, including 188 genes shared with the response to *G. ultimum*, 127 with *R. solani*, 112 with *A. brassicicola*, and 94 shared with both *A. brassicicola* and *R. solani*. Notably, 67 genes were consistently upregulated in response to contact with all tested conditions.

When considering differentially expressed genes with a fold change greater than 100, 59 genes were upregulated in response to *A. brassicicola* and 58 in response to *R. solani*, including 16 genes shared in the response to the two fungal pathogens. In contrast, only five genes were highly upregulated in response to the oomycete *G. ultimum*, none of which were shared with the fungal interactions. These results indicate that confrontation with fungal prey triggers a much stronger transcriptional activation in *T. atroviride* than interaction with the oomycete *G. ultimum*, in terms of intensity of induced genes. The most highly upregulated gene in response to *G. ultimum* showed a 771-fold increase after contact compared to before interaction, whereas in response to *A. brassicicola* and *R. solani*, 12 and 13 genes, respectively, exhibited fold changes exceeding 1,000.

Gene Ontology enrichment analyses were performed to identify the biological processes overrepresented within each category of differentially expressed genes. Upregulated genes in response to the two fungal pathogens, *A. brassicicola* and *R. solani*, were predominantly associated with the metabolism and catabolism of complex carbohydrates (Supplementary Figure 1). The most significantly enriched processes involved chitin degradation, aminoglycan catabolism, and glucosamine-containing compound metabolism. The coordinated activation of these catabolic pathways suggests a transcriptional response specifically directed toward the hydrolysis of fungal cell wall polymers and the utilization of associated amino sugars.

Genes specifically induced in response to *A. brassicicola* revealed a coordinated response structured around three major functional categories (Supplementary Figure 2). First, genes involved in lipid and sterol biosynthesis, particularly ergosterol and secondary alcohol biosynthetic pathways, were significantly enriched. Second, strong enrichment was observed for processes related to cell cycle progression, including mitosis, cell division, and cytokinesis. Third, genes associated with fungal cell wall organization were overrepresented. Together, these results indicate a transcriptional program oriented toward cellular proliferation and the structural reinforcement of membrane and cell wall compartments.

Genes specifically upregulated in response to *R. solani* were enriched in processes related to transmembrane transport, including the transport of monoatomic ions, cations, and carbohydrates (Supplementary Figure 3). Significant enrichment was also observed for pathways associated with fatty acid metabolism, particularly β-oxidation, as well as for processes involved in peroxisome organization and fungal cell wall biosynthesis, including chitin and aminoglycan synthesis. Consistent with these enrichments, seven of the 13 peroxisome biogenesis (PEX) genes identified in the *T. atroviride* genome were specifically upregulated in response to *R. solani* (PEX1, PEX2, PEX5, PEX11, PEX12, PEX13, and PEX14), whereas none were induced during interactions with *A. brassicicola* or *G. ultimum*. Among the genes specifically induced against *R. solani*, several were also involved in the shikimate pathway, which mediates the biosynthesis of aromatic amino acids. These included the DAHP synthase (ARO4), the pentafunctional AROM polypeptide (ARO1), and chorismate synthase (ARO2), which together catalyze the synthesis of chorismate, the common precursor of the three aromatic amino acids. In addition, ARO8, an aminotransferase involved in phenylalanine and tryptophan biosynthesis, was also upregulated in response to *R. solani*. Together, these results suggest a specific metabolic reprogramming of *T. atroviride* during interaction with *R. solani*, involving peroxisomal activity, lipid metabolism, and aromatic amino acid biosynthesis.

During confrontation with *G. ultimum*, upregulated genes were significantly enriched in pathways related to specialized metabolite biosynthesis and arabinan catabolism (Supplementary Figure 4). Similarly, genes shared between the responses to *A. brassicicola* and *G. ultimum* also displayed enrichment for specialized metabolite biosynthetic processes (Supplementary Figure 5). In contrast, no significantly enriched biological processes were detected among genes shared between *G. ultimum* and *R. solani*, nor among the genes commonly induced in response to all three pathogens. Unexpectedly, despite the identification of 395 genes specifically upregulated during self-confrontation between *T. atroviride* strains, no biological process was found to be significantly enriched within this gene set.

### 3.3 Pathogen-Specific Induction of Cell Wall-Degrading Enzymes and Effector-Like Proteins

Cell wall-degrading enzyme (CWDE) genes are among the gene families most strongly associated with mycoparasitism in *Trichoderma*. These genes encode enzymes involved in the degradation of cell wall polysaccharides, mostly glycoside hydrolases (GHs). GH genes were predominantly upregulated in response to both fungal pathogens (Figure 3A). Out of the 257 GH-encoding genes identified in the genome, 114 were upregulated in response to *A. brassicicola* and 112 in response to *R. solani*. During self-confrontation, 51 GH genes were upregulated in *T. atroviride*, of which 22 were shared with the responses to both fungal pathogens. In contrast, exposure to *G. ultimum* resulted in the upregulation of only 26 GH-encoding genes, 12 of which were also induced during self-confrontation.

**Figure 3:**
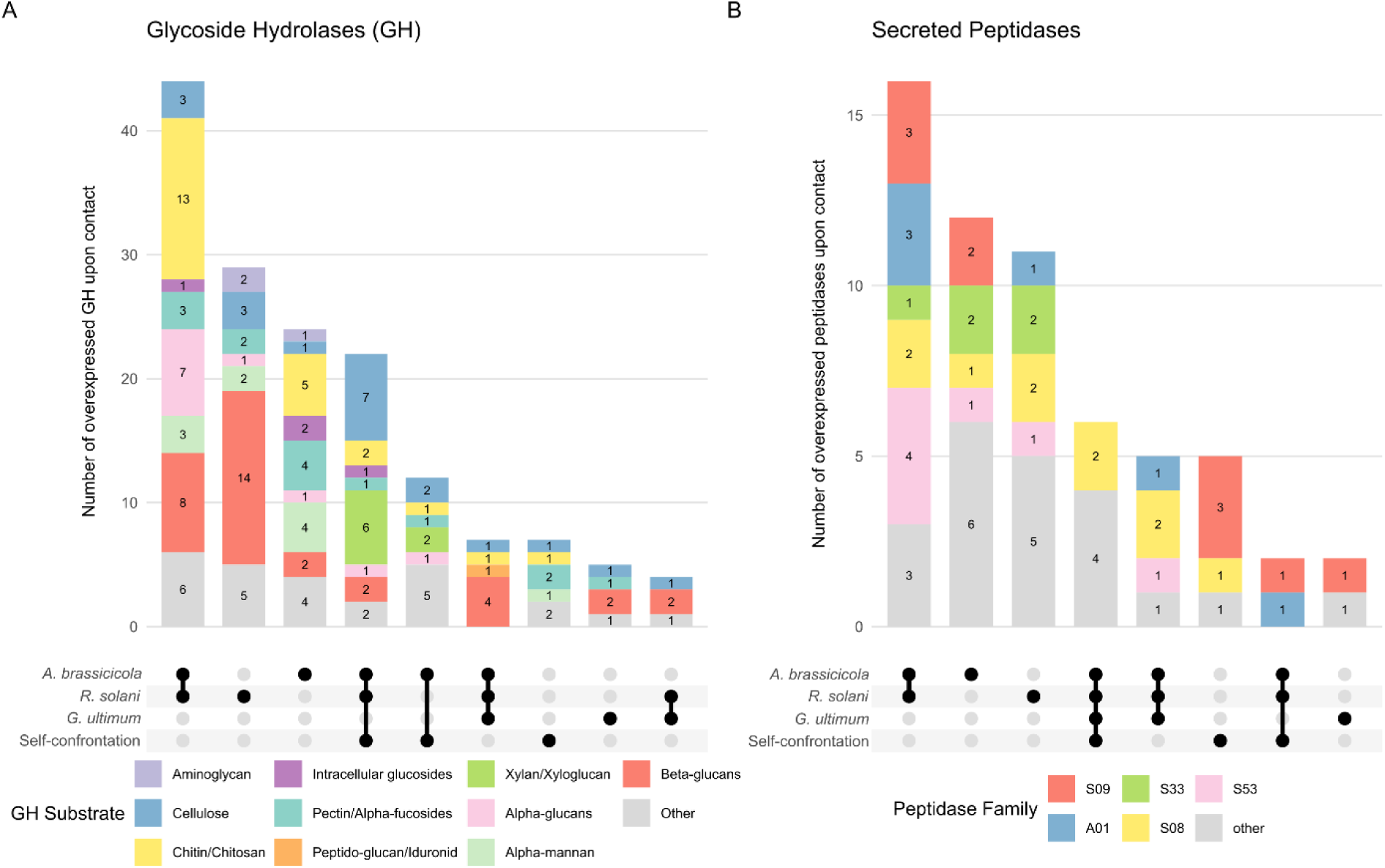
UpSet plot of CWDEs upregulated in response to contact. Genes upregulated in *Trichoderma atroviride* between before and after contact with *Alternaria brassicicola*, *Rhizoctonia solani*, *Globisporangium ultimum*, or during self-confrontation between *T. atroviride* strains. On the left (A), glycoside hydrolases (GHs), with colors representing the substrates degraded by these enzymes (“other” corresponding to β-mannans, α-galactosides, α-arabinofuranosides, β-xylosides, and α-glucuronosides). Conditions with fewer than 4 differentially expressed genes are not shown. On the right (B), secreted peptidases, with colors indicating the five most overrepresented peptidase families. Conditions with fewer than 2 differentially expressed genes are not shown.

Chitin is one of the most abundant polysaccharides found in fungal cell walls. Among the 41 genes encoding GHs involved in chitin and chitosan degradation, 13 were upregulated in response to contact with both *A. brassicicola* and *R. solani* (Figure 3A). These included eight chitinases (GH18), all belonging to class B chitinases, as well as four chitosanases (GH75) and one β-N-acetylglucosaminidase (GH20). Five additional genes were specifically induced in response to contact with *A. brassicicola*, including four chitinase-encoding genes (one class A, two class B, and one class C) and one GH20 gene. Two other chitinase genes were upregulated in response to both fungal prey and self-confrontation, including one class A and one class B chitinase. β-1,3-glucans are also major components of fungal cell walls, as well as those of oomycetes. Among the 47 GH genes involved in β-glucan degradation, four were upregulated in response to contact with all three pathogens (Figure 3A). Eight genes were upregulated during interactions with both *A. brassicicola* and *R. solani*, two were specifically induced in response to *A. brassicicola*, and 14 were specific to the interaction with *R. solani*. In contrast, only nine β-glucanase genes were upregulated in response to *G. ultimum*, compared with 16 and 31 during contact with *A. brassicicola* and *R. solani*, respectively.

Among the most highly upregulated genes in response to *A. brassicicola* (fold change > 100), 18 were associated with chitin degradation and four with β-glucan degradation. The most strongly induced gene during interaction with *A. brassicicola* encoded a β-glucanase, which was expressed 8,106-fold more after contact than before. In response to *R. solani*, four highly induced genes were involved in chitin degradation, including three belonging to the GH75 family, whereas six genes were associated with β-glucan degradation. In contrast, no CAZyme-encoding genes exhibited a fold change greater than 100 in response to *G. ultimum*.

Oomycete cell walls also contain cellulose, a polymer absent from fungal cell walls. However, among the 30 genes involved in cellulose degradation, only five were upregulated in response to contact with the oomycete *G. ultimum*, compared with 15, 16, and 12 genes upregulated during interactions with *A. brassicicola*, *R. solani*, and self-confrontation, respectively.

Secreted peptidase genes may also contribute to mycoparasitism by degrading structural proteins of the prey cell wall. Similar to GH genes, they were predominantly upregulated in response to both fungal pathogens (Figure 3B). Among the 120 secreted peptidases identified, 16 were specifically upregulated in response to both *A. brassicicola* and *R. solani*, while 12 were induced exclusively during interaction with *A. brassicicola* and 11 exclusively in response to *R. solani*. In addition, six secreted peptidase genes were upregulated under all tested conditions, and five were specifically induced in response to the three pathogens. Only two secreted peptidase genes were specifically upregulated in response to *G. ultimum*. The peptidases induced upon contact mainly belonged to the S class, particularly the S08, S09, S53, and S33 families, as well as the A01 family.

Some secreted peptidases were among the most highly upregulated genes in response to pathogen contact (fold change > 100). In interaction with *A. brassicicola*, two S08 family peptidases were strongly induced. During confrontation with *R. solani*, one M43 and one S53 peptidase were among the most highly expressed genes. In contrast, a single M35 peptidase was strongly induced in response to *G. ultimu*m and represented the most highly upregulated gene in this condition, with a fold change of 771.

Effectors, which are typically studied in the context of plant-pathogen interactions, may also play important roles in interactions between mycoparasitic fungi and their prey. These effectors are generally small secreted cysteine-rich proteins (SSCPs). Among the 182 genes encoding SSCPs identified in the genome, 23 were upregulated in response to both *A. brassicicola* and *R. solani* (Figure 4). These included proteins containing domains commonly associated with fungal effectors, such as two hydrophobins, two cerato-platanins, and one killer protein. An additional 23 SSCP-encoding genes were specifically induced in response to *R. solani*, including one cerato-platanin and one killer protein. In contrast, only nine SSCP-encoding genes were specifically upregulated in response to *A. brassicicola*, among which was a protein containing a CBM18 domain, a chitin-binding module potentially involved in fungal cell wall recognition. Exposure to *G. ultimum* induced a distinct set of SSCP-encoding genes, with nine genes specifically upregulated, including one encoding hydrophobin. Surprisingly, four genes encoding proteins containing a CBM1 domain were upregulated in response to *A. brassicicola*, *R. solani*, and self-confrontation with *T. atroviride*. Although CBM1 domains are involved in cellulose binding rather than cellulose hydrolysis, this observation is consistent with the concurrent upregulation of seven cellulase genes under the same conditions (Figure 3A).

**Figure 4:**
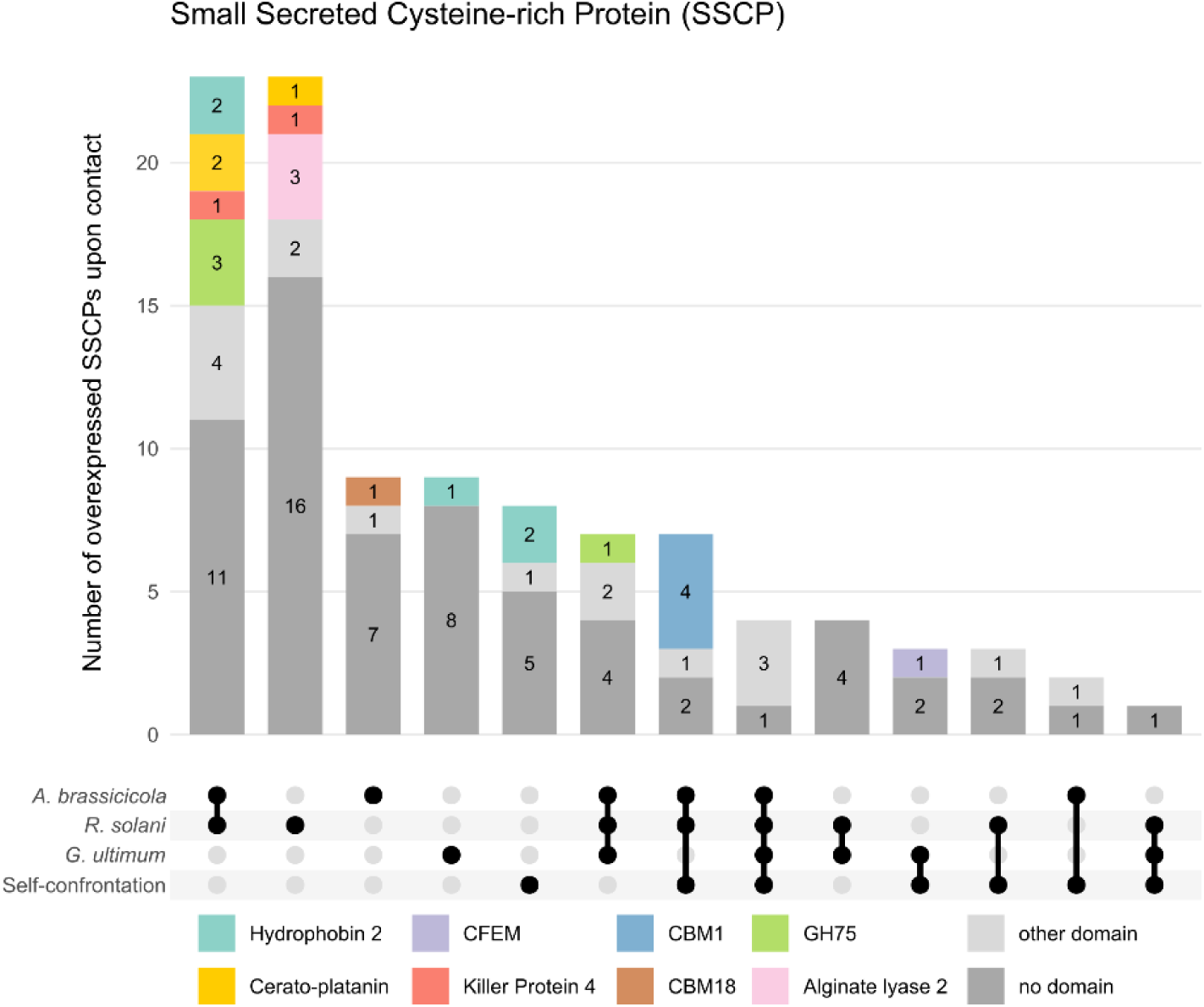
UpSet plot of Small Secreted Cysteine-rich Proteins upregulated in response to contact. Genes upregulated in *Trichoderma atroviride* between before and after contact with *Alternaria brassicicola*, *Rhizoctonia solani*, *Globisporangium ultimum*, or during self-confrontation between *T. atroviride* strains. SSCPs correspond to secreted proteins shorter than 300 amino acids and containing more than three cysteine residues. Colors indicate the PFAM domains identified within the proteins. CFEM = Common in Fungal Extracellular Membrane proteins; CBM = Carbohydrate-Binding Module; GH = Glycoside Hydrolase.

Among the most highly upregulated genes (fold change > 100) in response to *A. brassicicola*, 16 SSCP-encoding genes were identified, including two cerato-platanins. In response to *R. solani*, 21 SSCPs were among the most highly upregulated genes, seven of which were also highly induced in response to *A. brassicicola*, including one cerato-platanin.

Many GHs, secreted peptidases, and SSCPs are upregulated in response to pathogen contact in *T. atroviride*. This response is mainly observed against *A. brassicicola* and *R. solani*, compared to *G. ultimum*. Nevertheless, differences are still observed in the sets of upregulated genes between *A. brassicicola* and *R. solani*, although a substantial proportion is shared.

### 3.4 Pathogen-Specific Modulation of Specialized Metabolism

Specialized metabolite gene clusters were identified in *T. atroviride* genome. Expression of the 57 biosynthetic genes contained within the 46 predicted clusters was assessed between before and after contact with the different pathogens.

Figure 5 shows that a large number of biosynthetic genes were upregulated in response to pathogen contact, whereas relatively few were induced during self-confrontation. Marked differences were also observed among the three pathogens, with only three biosynthetic genes commonly upregulated in response to all three. In response to *A. brassicicola* and *G. ultimum*, 17 biosynthetic genes were upregulated, compared to 14 in response to *R. solani*. Eight genes were shared between *R. solani* and *A. brassicicola*, five between *G. ultimum* and *R. solani*, and seven between *G. ultimum* and *A. brassicicola*. All major classes of specialized metabolite biosynthetic gene clusters were represented in the responses to the three pathogens, including NRPS (Non-Ribosomal Peptide Synthetase), PKS (Polyketide Synthases), terpene synthases, and isocyanide synthases. Numerous biosynthetic genes show pathogen-specific expression patterns, particularly in response to *A. brassicicola* and *G. ultimum*, where five and seven biosynthetic genes are upregulated, respectively.

**Figure 5:**
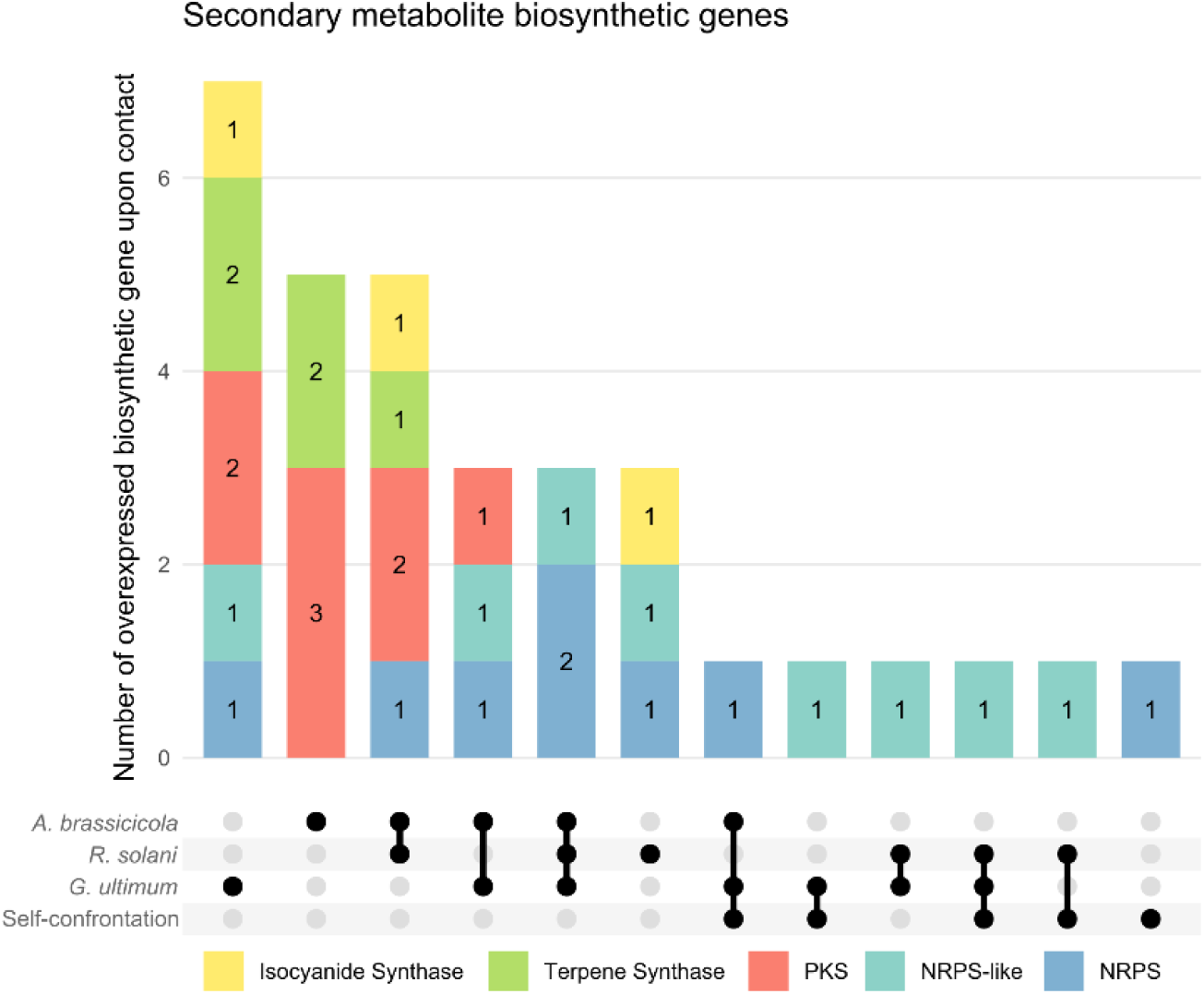
UpSet plot of specialized metabolite biosynthetic genes upregulated in response to contact. Genes upregulated in *Trichoderma atroviride* between before and after contact with *Alternaria brassicicola*, *Rhizoctonia solani*, *Globisporangium ultimum*, or during self-confrontation between *T. atroviride* strains. Forty-six specialized metabolite biosynthetic gene clusters were identified in *T. atroviride*, and the 57 biosynthetic genes contained within these clusters were analyzed. PKS = Polyketide Synthase, NRPS = Non-Ribosomal Peptide Synthetase.

Among the most highly upregulated genes in *T. atroviride* during pathogen interaction (fold change > 100), genes involved in specialized metabolite biosynthesis were exclusively detected in response to *A. brassicicola*. These included five biosynthetic genes distributed across three biosynthetic gene clusters: a PKS and an NRPS-like gene located within the same cluster, a PKS gene associated with a second cluster, and an NRPS together with an NRPS-like gene within a third cluster. This latter cluster comprises four biosynthetic genes, with the NRPS and NRPS-like genes showing fold changes above 100, while the two additional PKS genes in the same cluster also displayed strong induction levels (fold changes of 19 and 68) in response to *A. brassicicola*. The genes within this cluster were also more highly expressed in highly parasitic *T. atroviride* strains than in weakly parasitic strains (Brémand et al., 2026a).

### 3.5 Pathogen-Specific Perception by *T. atroviride*

Analysis of genes whose expression is upregulated in response to *G. ultimum* revealed that a large proportion of them encode proteins containing NACHT domains. These domains are characteristic of NLR (NOD-like receptor) proteins, which are intracellular receptors known to be involved in immunity (Uehling et al., 2017). In fungi, NLR proteins mainly contain either NACHT or NB-ARC domains as their central nucleotide-binding domain (Bonometti et al., 2025).

In the genome of *T. atroviride*, 163 potential fungal NLRs were identified: 125 contained a NACHT domain (77%), 28 contained an NB-ARC domain (17%), one possessed both domains, and nine lacked both NACHT and NB-ARC domains but contained N-terminal domains commonly associated with fungal NLRs, such as NACHT_N, Helo-like, or PNP_UDP. The most frequent N-terminal domains identified in *T. atroviride* NLRs were PNP_UDP (32%), NACHT_N (15%), HET (12%), and Helo-like (11%), with some proteins containing multiple domains. Regarding C-terminal domains, ankyrin repeats were the most abundant (41%), followed by WD40 repeats (25%) and TPR domains (17%).

Among these putative fungal NLR-encoding genes, 42 were upregulated in response to *A. brassicicola*, 7 in response to *R. solani*, 63 in response to *G. ultimum*, and 33 in response to self-confrontation (Figure 6A). The expression of these genes therefore appeared to be more strongly associated with responses to *A. brassicicola* and *G. ultimum* than to *R. solani*. However, the sets of induced genes differed substantially between conditions, with only 18 shared between the responses to *A. brassicicola* and *G. ultimum*. Notably, 25 putative fungal NLR-encoding genes were specifically upregulated in response to *G. ultimum*, including a relatively high proportion of NB-ARC-containing proteins compared with the other conditions. Another 25 NLRs were shared between the responses to *G. ultimum* and self-confrontation.

**Figure 6:**
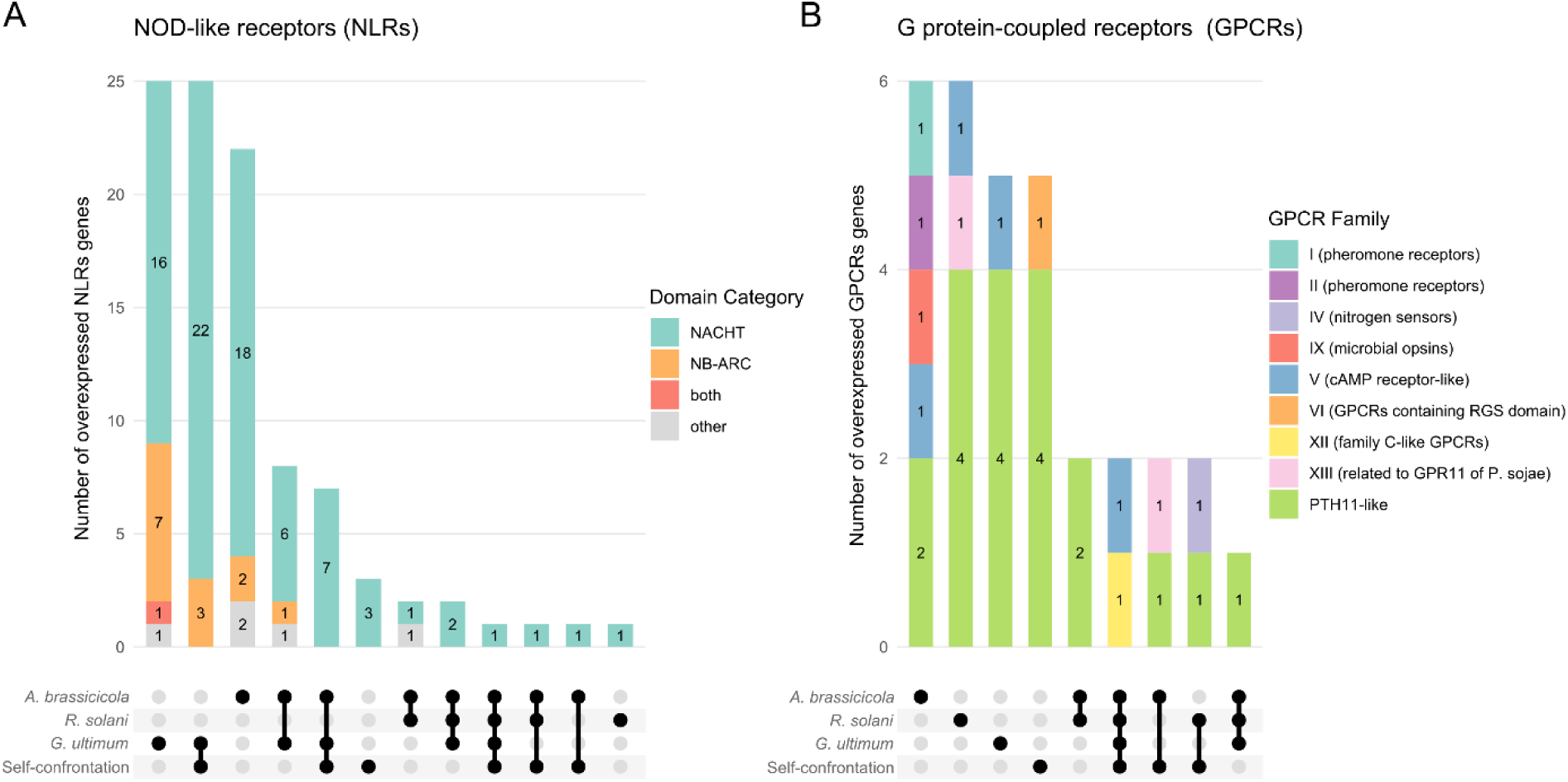
UpSet plot of receptor-encoding genes upregulated in response to contact. Genes upregulated in *Trichoderma atroviride* between before and after contact with *Alternaria brassicicola*, *Rhizoctonia solani*, *Globisporangium ultimum*, or during self-confrontation between *T. atroviride* strains. Left panel shows genes encoding NLRs (NOD-like receptors), categorized according to their central domain (A). Right panel shows genes encoding GPCRs (G protein-coupled receptors) identified by Gruber et al. (2013) (B).

In response to *A. brassicicola*, 22 NLR-encoding genes were specifically upregulated (Figure 6A). Surprisingly, only two NLRs were shared between the responses to both fungal pathogens, and only one was specifically induced in response to *R. solani*. In contrast to other mechanisms such as CWDEs and SSCPs, which showed a strong common response to both fungal pathogens, NLR expression patterns differed markedly between *A. brassicicola* and *R. solani*. For these genes, the response to *A. brassicicola* appeared more similar to that observed against *G. ultimum*, although the sets of induced genes remained largely distinct, with only six genes specifically shared in response to these two pathogens.

However, no study has yet demonstrated the involvement of NLRs in prey perception by *Trichoderma*. Another class of receptors that has been much more extensively studied in *Trichoderma* is the G protein-coupled receptors (GPCRs), which are membrane-bound receptors involved in signal transduction (Omann and Zeilinger, 2010). All 65 GPCRs previously identified in *T. atroviride* by Gruber et al. (2013) were recovered in our analysis through sequence similarity searches.

Regarding their expression, 13 GPCR-encoding genes were upregulated during interactions with *A. brassicicola* and *R. solani*, with five genes shared between both interactions (Figure 6B). In contrast, only eight GPCR-encoding genes were induced during interaction with *G. ultimum*, including three that were also responsive to the fungal pathogens. A large proportion of GPCR-encoding genes exhibited condition-specific expression patterns. Six genes were exclusively upregulated during interaction with *A. brassicicola*, six during interaction with *R. solani*, five during interaction with *G. ultimum*, and five during self-confrontation. Most of these genes encoded PTH11-like GPCRs. Notably, the interaction with *A. brassicicola* was associated with a greater diversity of GPCR classes, including receptors potentially involved in pheromone sensing and cAMP-mediated signaling pathways.

The differential expression of genes encoding these two types of receptors suggests that *T. atroviride* may differentially perceive the three pathogens, potentially reflecting pathogen-specific recognition. Such differences in pathogen perception could contribute to the distinct mycoparasitic mechanisms deployed against each pathogen and may help explain the pathogen specificity observed in the antagonistic response.

## 4 Discussion

Species belonging to the genus *Trichoderma* exhibit a broad spectrum of antagonistic activity and can be used against a wide range of phytopathogens. However, not all *Trichoderma* strains are equally effective against all target organisms. Our previous studies have highlighted substantial variability in antagonistic performance both between and within *Trichoderma* species (Brémand et al., 2026b), with marked interspecific differences in biocontrol efficiency and pronounced intraspecific variation among the tested *T. atroviride* strains. In agreement with these observations, distinct antagonistic mechanisms have been described among *Trichoderma* species (Atanasova et al., 2013), but also between highly and weakly mycoparasitic strains of *T. atroviride* (Brémand et al., 2026a). Differences in host specificity have also been observed according to the phylogenetic clade of the antagonist. Strains belonging to the *Longibrachiatum* clade were generally more effective against the oomycete *G. ultimum*, whereas strains from the *Viride* clade displayed stronger antagonistic activity against the fungal pathogens *A. brassicicola* and *R. solani* (Brémand et al., 2026b). These differences in host range may be explained by the distinct mechanisms employed by strains from these clades in response to a given pathogen (Atanasova et al., 2013). These observations suggest that the outcome of *Trichoderma*-pathogen interactions depends on both the identity of the antagonist and the target pathogen, and is shaped by the molecular mechanisms deployed by *Trichoderma*. In the present study, the four *T. atroviride* strains were specifically selected for their strong antagonistic activity against the three tested pathogens (Figure 1A). Consequently, only limited differences in antagonistic intensity were observed between pathogen species. Nevertheless, including multiple strains for each pathogen species revealed significant intraspecific variation in susceptibility to *T. atroviride*. Such strain-dependent differences were particularly evident for *R. solani* and *G. ultimum* (Figure 1B). This variability could lead to differences in the efficiency of *Trichoderma*-based biocontrol products under field conditions, as the composition of pathogen populations may vary across geographical regions and growing seasons. Moreover, the repeated use of *T. atroviride* raises questions about the potential emergence of pathogen resistance to its antagonistic mechanisms and, consequently, the long-term durability of this biocontrol strategy (Bardin et al., 2015).

Despite the limited phenotypic variation observed among *T. atroviride* strains in their antagonistic activity against the three pathogens, substantial transcriptomic divergence was detected. Among the 3,737 genes differentially expressed between pre-contact and post-contact conditions in at least one of the tested conditions, only 2% were shared across the responses to all three pathogens (Figure 2). The transcriptional responses to *A. brassicicola* and *R. solani* were more closely related, sharing 11% of differentially expressed genes. In contrast, the overlap involving *G. ultimum* was markedly lower. These results reveal a clear divergence between the responses of *T. atroviride* to fungal and oomycete pathogens. This observation is consistent with our previous study reporting a strong correlation in the efficiency of diverse *Trichoderma* strains against *A. brassicicola* and *R. solani*, but a much weaker correlation against *G. ultimum* (Brémand et al., 2026b). Together, these findings indicate that *Trichoderma* strains do not rely on the same molecular mechanisms against all plant pathogens. In this context, broadly effective strains appear to combine multiple antagonistic strategies, enabling them to maintain activity against both fungal and oomycete pathogens through the deployment of distinct and complementary molecular responses. Most of the upregulated genes were pathogen-specific, with 23% uniquely induced in response to *A. brassicicola*, 11% specific to *R. solani*, and 11% specific to *G. ultimum* (Figure 2). Notably, even between the two fungal pathogens, *A. brassicicola* and *R. solani*, *T. atroviride* activated distinct transcriptional programs, highlighting the high degree of specialization underlying its antagonistic response. Indeed, expression of mycoparasitism-related genes was monitored by RT-qPCR in *T. harzianum* during confrontation with *Fusarium oxysporum*, *Colletotrichum capsici*, *Colletotrichum truncatum*, and *Gloeocercospora sorghi*, revealing strong host-dependent variations in transcriptional responses (Sharma et al., 2017).

The genes commonly upregulated in *T. atroviride* during interaction with both fungal pathogens are predominantly associated with attack-related mechanisms. Gene Ontology enrichment analyses most prominently highlighted the overrepresentation of genes involved in polysaccharide degradation, particularly GH. These enzymes include key actors in chitin degradation (chitinases, chitosanases, and β-N-acetylglucosaminidases) as well as β-1,3-glucanases involved in the breakdown of fungal cell walls. Among these genes, eight chitinases were identified, all belonging to class B. This observation supports the hypothesis that class B chitinases are more specifically involved in mycoparasitism than classes A and C. This hypothesis is consistent with previous genomic analyses showing an enrichment of class B chitinases in *Trichoderma* compared with other fungi, as well as in mycoparasitic species such as *T. atroviride* and *T. virens* relative to the saprotrophic species *T. reesei* (Seidl et al., 2005; Wang et al., 2023a). However, some GH-encoding genes were specifically induced depending on the pathogen encountered (Figure 3A). The response to *A. brassicicola* involved a stronger induction of genes involved in chitin degradation, whereas interaction with *R. solani* preferentially activated genes associated with β-glucan degradation. Consistent with this, Atanasova et al. (2013) reported that *T. atroviride* expresses a higher number of β-glucanase encoding genes during interaction with *R. solani* compared to *T. virens* and *T. reesei*. Similarly, Enriquez-Felix et al. (2024) observed differential CAZyme-encoding genes expression in *T. atroviride* during interactions with *Alternaria alternata* and *R. solani*, with approximately twice as many CAZyme-encoding genes induced against *A. alternata*. In our study, a comparable number of GH-encoding genes were upregulated in response to *A. brassicicola* and *R. solani* (106 genes each), with a substantial overlap of 71% (75 shared genes), highlighting both a conserved core response and a significant degree of pathogen-specific transcriptional regulation.

The degradation of host cell walls also requires the secretion of peptidases involved in the hydrolysis of structural proteins. Genes encoding these enzymes were predominantly induced upon contact with *A. brassicicola* and *R. solani* (Figure 3B). Among the 120 genes, 14 were specifically upregulated in response to both pathogens, while 11 and 10 were uniquely induced by *A. brassicicola* and *R. solani*, respectively. Peptidases have previously been suggested to play a central role in *Trichoderma* mycoparasitism. In *T. harzianum*, the top 10 most highly induced genes during interaction with *Sclerotinia sclerotiorum* contains predominantly peptidase-encoding genes rather than GHs (Steindorff et al., 2014). This trend appears even more pronounced in *T. atroviride*, which shows a stronger induction of peptidase genes during interaction with *R. solani* compared to *T. virens* and *T. reesei* (Atanasova et al., 2013). At the genomic level, peptidase families, particularly S8 serine peptidase, are among the most expanded in *Trichoderma* compared to other fungi. While the average copy number of S8 peptidase-genes is approximately 9.6 in fungi and 10 in the saprotrophic species *T. reesei*, it increases substantially in mycoparasitic species, reaching 33 in *T. virens* and 36 in *T. atroviride* (Kubicek et al., 2011). Together, these observations suggest that peptidase secretion is a major component of the mycoparasitic arsenal in *Trichoderma*, and particularly in *T. atroviride*, where it appears to be associated with efficient mycoparasitic performance (Brémand et al., 2026a).

Another mechanism that appeared to be preferentially associated with the response to the two fungal pathogens is the production of small secreted cysteine-rich proteins (SSCPs). Of the 182 predicted SSCP-encoding genes identified in the *T. atroviride* genome, 52 and 72 were upregulated in response to *A. brassicicola* and *R. solani*, respectively (Figure 4). Several of these proteins contained domains commonly found in fungal effectors, including hydrophobins, cerato-platanins, and killer proteins. The importance of SSCPs during mycoparasitism has previously been highlighted by Atanasova et al. (2013), who reported a stronger induction of genes encoding effector-like proteins in *T. atroviride* than in *T. virens* and *T. reesei* during interaction with *R. solani*. Functional characterization of several SSCPs has further demonstrated their contribution to antagonistic interactions. For example, the LysM effector Tal6 was shown to participate in both mycoparasitism and plant association in *T. atroviride* (Romero-Contreras et al., 2019), while a hydrophobin from *T. virens* enhances antagonistic activity against *R. solani* and promotes root colonization of *Arabidopsis thaliana* (Guzmán-Guzmán et al., 2017). Furthermore, substantial variation was observed between the responses to *A. brassicicola* and *R. solani*, with each pathogen inducing a distinct subset of SSCPs, particularly in the case of *R. solani*. Similar pathogen-specific expression patterns have previously been reported in *T. atroviride*, including a cerato-platanin specifically induced during interaction with *R. solani* and a CFEM domain-containing protein specifically induced in response to *Alternaria alternata* (Enriquez-Felix et al., 2024).

Unexpectedly, relatively few secreted GH-, peptidase- and SSCP-encoding genes were upregulated in response to the oomycete *G. ultimum* (Figure 3). Only 26 GH-, 16 peptidase-, and 28 SSCP-encoding genes were induced, which is substantially lower than the responses observed against the fungal pathogens *A. brassicicola* (114 GHs, 43 peptidases and 52 SSCPs) and *R. solani* (112 GHs, 43 peptidases and 73 SSCPs). However, despite this reduced diversity, the GH-encoding genes induced in response to *G. ultimum* showed a distinct functional bias. Indeed, nine GHs were involved in β-glucan degradation and five in cellulose degradation, representing a higher relative proportion of cellulose-degrading enzymes (19%) compared with *A. brassicicola* (13%) and *R. solani* (14%), as well as an increased proportion of β-glucanases against *G. ultimum* (35%) compared with *A. brassicicola* (14%) and *R. solani* (28%). Several studies have similarly highlighted the importance of cellulases and β-glucanases during interactions between *Trichoderma* and oomycetes. Wang et al. (2024) reported the induction of CWDEs in *T. harzianum* during interaction with *Phytophthora capsici*. In addition, cellulase and peptidase production has been observed in six *Trichoderma* species, including *T. atroviride*, in response to *Pythium myriotylum* mycelium, with cellulase activity correlating positively with strain aggressiveness (Chen et al., 2024). Interestingly, exposure to inactivated *P. myriotylum* mycelium was also shown to induce β-1,3-glucanases and cellulases in *T. harzianum*. Moreover, disruption of the cellulase-associated transcription factor XYR1 had only a limited impact on mycelial degradation capacity in *T. harzianum*, suggesting a major contribution of β-1,3-glucanases in the interaction with *P. myriotylum* (Anjago et al., 2026). In contrast, in *T. reesei*, XYR1 deletion resulted in reduced antagonistic activity against *Pythium* and *Globisporangium* (Chen et al., 2024), but cellulase production remained the primary induced response during fungal prey interaction in this species (Atanasova et al., 2013). Together, these observations highlight the importance of cellulases and β-glucanases in *Trichoderma* interactions with oomycetes. Consistently, both enzyme classes were also identified in our dataset, with genes encoding five cellulases and nine β-glucanases specifically induced in response to *G. ultimum*. However, these numbers remained lower than those observed in interactions with fungal pathogens, where 15 and 16 cellulase-encoding genes and 16 and 31 β-glucanase-encoding genes were induced in response to *A. brassicicola* and *R. solani*, respectively. The results also identified several secreted peptidases and SSCPs that were specifically upregulated in response to *G. ultimum* and may contribute to this interaction. Notably, the most highly upregulated gene in response to *G. ultimum* encodes a secreted M35 peptidase, which may play a role in the antagonistic response to this pathogen.

All of these attack-related mechanisms appeared to be less prominently induced during interaction with *G. ultimum* compared with the two fungal pathogens. Instead, Gene Ontology enrichment analysis of genes upregulated in response to *G. ultimum* revealed a significant overrepresentation of functions associated with specialized metabolite biosynthesis. Consistently, *T. atroviride* upregulated 17 specialized metabolite biosynthetic genes in response to *G. ultimum* (Figure 5). A similar number was upregulated during interaction with *A. brassicicola* (17 genes), whereas 14 genes were induced in response to *R. solani*. However, despite these comparable numbers, the identity of the induced genes varied markedly between pathogens. Indeed, five, three, and seven biosynthetic genes were specifically upregulated in response to *A. brassicicola*, *R. solani*, and *G. ultimum*, respectively, highlighting a strong pathogen-dependent reprogramming of specialized metabolism. Atanasova et al. (2013) showed that *T. atroviride* activates a broad repertoire of specialized metabolite biosynthetic genes during interaction with *R. solani*, whereas *T. virens* mainly induced genes involved in gliotoxin biosynthesis and *T. reesei* displays only limited induction of specialized metabolism, highlighting the importance of this mechanism in *T. atroviride*. Enriquez-Félix et al. (2024) reported pathogen-specific expression of biosynthetic genes in *T. atroviride*, including PKS and NRPS genes induced during interaction with *R. solani* and distinct NRPS genes induced in response to *A. alternata*. This transcriptional plasticity may have functional significance, as the activity spectrum of specialized metabolites is often restricted to specific targets. A well-known example is provided by two strains of *T. virens*, where one produces gliovirin and the other gliotoxin: the first is effective against *G. ultimum* but not *R. solani*, whereas the second shows the opposite pattern (Howell et al., 1993). Together, these observations suggest that modulation of specialized metabolite biosynthesis represents a key adaptive component of the *T. atroviride* response, enabling the deployment of distinct chemical arsenals depending on the nature of the pathogen encountered.

Among the pathogen-specific responses identified in *T. atroviride*, interaction with *A. brassicicola* and *R. solani* was characterized by a strong overrepresentation of genes associated with cell wall remodeling. These transcriptional changes suggest that extensive structural and physiological adaptations occur during confrontation with these pathogens. Previous studies have highlighted the central role of cell wall remodeling during mycoparasitism in *T. atroviride*. Kappel et al. (2020) reported that several genes involved in chitin synthesis and deacetylation are induced under cell wall and oxidative stress conditions, including NaCl-, H₂O₂-, and SDS-induced stress, suggesting that these pathways contribute to stress adaptation and cell wall plasticity. The same study further demonstrated that the expression of these genes varies depending on the target pathogen, with stronger induction observed during interaction with *S. sclerotiorum* compared with *B. cinerea* or *R. solani*. Importantly, mutants impaired in these pathways displayed severe defects in overgrowing of *S. sclerotiorum* and *B. cinerea*. In our study, five of the chitin synthases identified by Kappel et al. (2020) were upregulated in response to *R. solani*. More recently, Kappel et al. (2024) showed that the cell wall of *T. atroviride* becomes enriched in chitosan upon contact with *S. sclerotiorum*. These findings indicate that dynamic regulation of cell wall architecture constitutes a key adaptive mechanism during antagonistic interactions with fungal pathogens. In contrast, much less information is currently available regarding the role of membrane remodeling during mycoparasitism in *Trichoderma*. Nevertheless, the overrepresentation of genes associated with lipid metabolism and membrane organization observed against *A. brassicicola* suggests that membrane dynamics may also constitute an important component of the antagonistic response.

Interaction with *R. solani* was characterized by the specific induction of genes associated with fatty acid metabolism, β-oxidation, and peroxisome organization. Notably, seven of the 13 peroxisome biogenesis (PEX) genes identified in the *T. atroviride* genome were specifically upregulated during interaction with *R. solani*. This coordinated induction of PEX genes suggests an important reorganization of peroxisomal functions during confrontation with this pathogen. Peroxisomes are central organelles involved in fatty acid β-oxidation, reactive oxygen species detoxification, and lipid metabolism, processes that are closely linked to fungal development and stress adaptation (Maruyama and Kitamoto, 2013). Interestingly, in our study, this mechanism appeared to be specifically induced in response to the basidiomycete *R. solani*, but not during interaction with the ascomycete *A. brassicicola* or the oomycete *G. ultimum*. Consistent with this observation, to our knowledge, no previous RNA-seq studies investigating interactions between *T. atroviride* and ascomycete fungal pathogens have reported induction of peroxisome-associated pathways. In contrast, induction of peroxisome-related genes has previously been described during interaction between *T. atroviride* and the basidiomycete *Armillaria ostoyae* (Chen et al., 2023), suggesting that activation of these pathways may be associated with antagonistic responses toward specific groups of fungal pathogens. Surprisingly, induction of peroxisome-associated genes has also been observed during interaction between *T. asperellum* and the pine wood nematode (Chen et al., 2024). In addition, comparative analyses between strongly and weakly mycoparasitic *T. atroviride* strains revealed a higher expression of peroxisome- and β-oxidation-related genes in highly parasitic strains, further supporting a link between peroxisomal metabolism and mycoparasitic performance (Brémand et al., 2026a). Together, these observations suggest that peroxisome remodeling and lipid catabolism may represent important adaptive mechanisms underlying efficient antagonistic activity in *T. atroviride*, particularly during interactions with basidiomycete pathogens.

The genes specifically induced during interaction with *R. solani* included genes involved in the shikimate pathway. This pathway is responsible for the biosynthesis of chorismate, the common precursor of the aromatic amino acids phenylalanine, tyrosine, and tryptophan, suggesting an activation of aromatic amino acid metabolism during confrontation with *R. solani*. Atanasova et al. (2013) observed the overexpression of genes associated with aromatic amino acid biosynthesis in *T. virens* during overgrowth of *R. solani*, notably the gene encoding DAHP synthase, the first and regulatory enzyme of the shikimate pathway. The authors suggested that activation of this pathway may reflect an increased demand for aromatic amino acids and potentially contribute to the biosynthesis of specialized metabolites involved in antagonism. Additional evidence supporting the importance of this pathway in *Trichoderma* biocontrol was provided by Pérez et al. (2015), who characterized the chorismate mutase gene in *T. parareesei*. Silencing of this gene resulted in reduced growth, impaired mycoparasitic activity against *R. solani*, *F. oxysporum*, and *B. cinerea*, as well as diminished root colonization ability. These findings highlight the role of the shikimate pathway not only in fungal metabolism, but also in the regulation of antagonistic activity.

To activate pathogen-responsive genes, *Trichoderma* must be able to perceive its interacting organism and initiate appropriate downstream signaling pathways. The strong transcriptional plasticity observed in our study suggests that these sensing systems may discriminate among different pathogens and contribute to the deployment of pathogen-specific responses. GPCRs are among the best-characterized receptor classes in *Trichoderma* and act as membrane-localized sensors of external signals, including host-derived molecules and cell wall degradation products (Omann and Zeilinger, 2010). Among the GPCR-encoding genes identified by Gruber et al. (2013) in *T. atroviride*, 13 and 14 were upregulated in response to *A. brassicicola* and *R. solani*, respectively, with five genes shared between both interactions (Figure 6B). Eight GPCRs were also induced during interaction with *G. ultimum*. Given that certain GPCRs have been shown to regulate the expression of genes encoding CWDEs in *Trichoderma* (Omann and Zeilinger, 2010), some of the GPCRs induced in our study may contribute to pathogen recognition and the subsequent activation of antagonistic mechanisms. The overlap in GPCR induction between *A. brassicicola* and *R. solani* is consistent with the activation of shared mycoparasitic responses, including CWDE production, whereas pathogen-specific GPCRs may contribute to the differential regulation of these responses. NLRs showed a distinct expression pattern. NLR-encoding genes were predominantly upregulated in response to *A. brassicicola* and *G. ultimum*, but with very limited overlap between the two conditions (Figure 6A). This suggests that distinct subsets of NLRs may be recruited depending on the interacting pathogen and could contribute to pathogen-specific transcriptional reprogramming. This hypothesis is consistent with the high specificity reported for fungal NLRs (Uehling et al., 2017), in contrast to GPCRs, which are generally considered broader environmental sensors (Omann and Zeilinger, 2010).

The near absence of NLR induction during interaction with the basidiomycete *R. solani*, compared to the two other pathogens, may also reflect the evolutionary history of mycoparasitism in *Trichoderma*. Early-diverging mycoparasitic relatives such as *Hypomyces*, *Sphaerostilbella*, *Arachnocrea*, and *Escovopsis* are largely specialized on basidiomycete hosts, suggesting that ancestral *Trichoderma*-like fungi may have primarily targeted this group. The ability of *Trichoderma* species to efficiently parasitize phytopathogenic ascomycetes likely represents a more recent evolutionary innovation (Chaverri and Samuels, 2013). Phylogenetic and ecological analyses further indicate that the evolution of *Trichoderma* involved multiple host jumps and transitions toward ecological niches associated with plant-pathogenic ascomycetes. In addition, Druzhinina et al. (2018) proposed a molecular basis for this adaptation, showing that *Trichoderma* acquired numerous lytic enzymes through horizontal gene transfer from plant-associated ascomycete fungi, thereby increasing its trophic versatility and ability to exploit phylogenetically diverse hosts. In this context, the expression patterns observed in this study raise the possibility that NLRs are more specifically involved in the recognition of recently acquired hosts such as ascomycetes and oomycetes, whereas the perception of basidiomycetes may rely more heavily on conserved sensing systems mediated by GPCRs, reflecting ancestral mycoparasitic strategies. This hypothesis is further supported by the marked expansion of NLR-coding genes in the genus *Trichoderma*, and particularly in the mycoparasitic species *T. atroviride* and *T. virens* compared with the saprotrophic species *T. reesei* (Dyrka et al., 2014).

## 5 Conclusion

This study provides a comprehensive transcriptomic overview of *Trichoderma atroviride* responses during interaction with three phylogenetically distinct phytopathogens, revealing a high degree of transcriptional plasticity underlying its antagonistic lifestyle. Despite relatively limited phenotypic variation in mycoparasitic performance among strains, RNA-seq analyses uncovered extensive and highly structured transcriptional reprogramming, with a large fraction of differentially expressed genes being pathogen-specific and only a limited core response shared across all interactions.

Our results support a model in which *T. atroviride* does not rely on a single conserved antagonistic program, but instead integrates multiple layers of perception and response to tailor its molecular strategy to the identity of the target organism. In this framework, interactions with the two fungal pathogens, *A. brassicicola* and *R. solani*, were characterized by partially overlapping yet clearly differentiated responses involving CWDEs, SSCPs, and specialized metabolism (Figure 7). In contrast, the response to the oomycete *G. ultimum* was markedly distinct, with reduced engagement of classical mycoparasitic functions and a shift toward alternative specialized metabolite biosynthetic programs.

**Figure 7.**
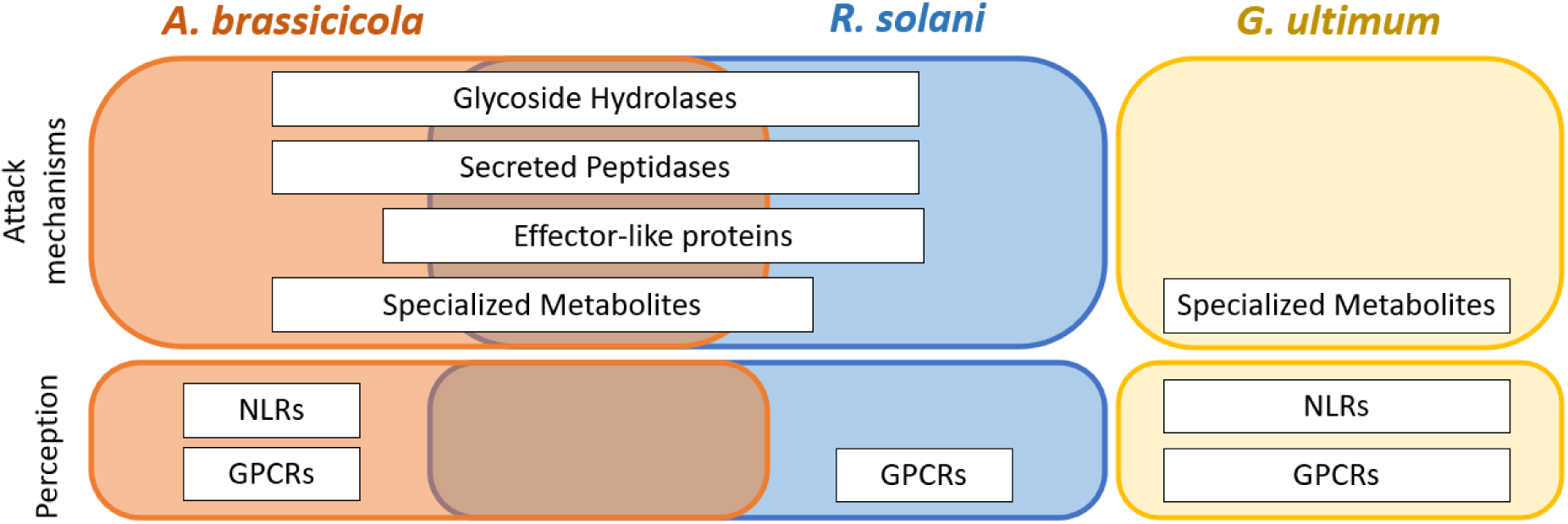
Schematic representation of the perception and attack mechanisms deployed by *T. atroviride* in response to the three pathogens. Mechanisms specifically induced in response to *Alternaria brassicicola* are shown in orange, those specific to *Rhizoctonia* solani in blue, and those specific to *Globisporangium ultimum* in yellow. Mechanisms shared between the responses to *A. brassicicola* and *R. solani* are represented in the overlapping region between the orange and blue areas.

Alongside this plasticity in attack mechanisms, we also observed plasticity in pathogen perception, with GPCRs and NLRs largely overexpressed in response to a single pathogen (Figure 7). This suggests that transcriptional plasticity may be regulated through fine-tuned perception of different prey. Together, these findings indicate that antagonism in *T. atroviride* relies on the coordinated regulation of pathogen perception and downstream attack mechanisms, enabling distinct molecular responses according to the target organism. Overall, our results support a model in which the antagonistic versatility of *T. atroviride* emerges from the dynamic coordination of conserved molecular functions rather than from a single universal response.

## Acknowledgments

We would like to thank the University of Angers and Angers Loire Métropole for funding part of the PhD work associated with this publication. We are grateful to the ANAN platform (SFR QUASAV), especially Muriel Bahut, and to the PACEM platform (SFR ICAT), especially Jérôme Cayon, for technical and material support. We also acknowledge the GenOuest bioinformatics core facility (https://www.genouest.org) for providing computational infrastructure.

## Data availability

All raw RNA sequencing data are accessible at the National Center for Biotechnology Information (NCBI) under BioProject accession PRJEB108439 (ncbi.nlm.nih.gov/bioproject/PRJEB108439). A comprehensive list of all gene identifiers and annotations discussed in this study is provided in Supplementary Dataset 1. All scripts used are available on GitHub (github.com/ebremand/Trichoderma-Transcriptomic-Plasticity) and have been deposited on Zenodo (doi.org/10.5281/zenodo.21070911).

## Notes

### Competing Interest Statement

The authors have declared no competing interest.

