## Supplementary Figures for "Transcriptional Plasticity of *Trichoderma atroviride* in Response to Different Plant Pathogens"

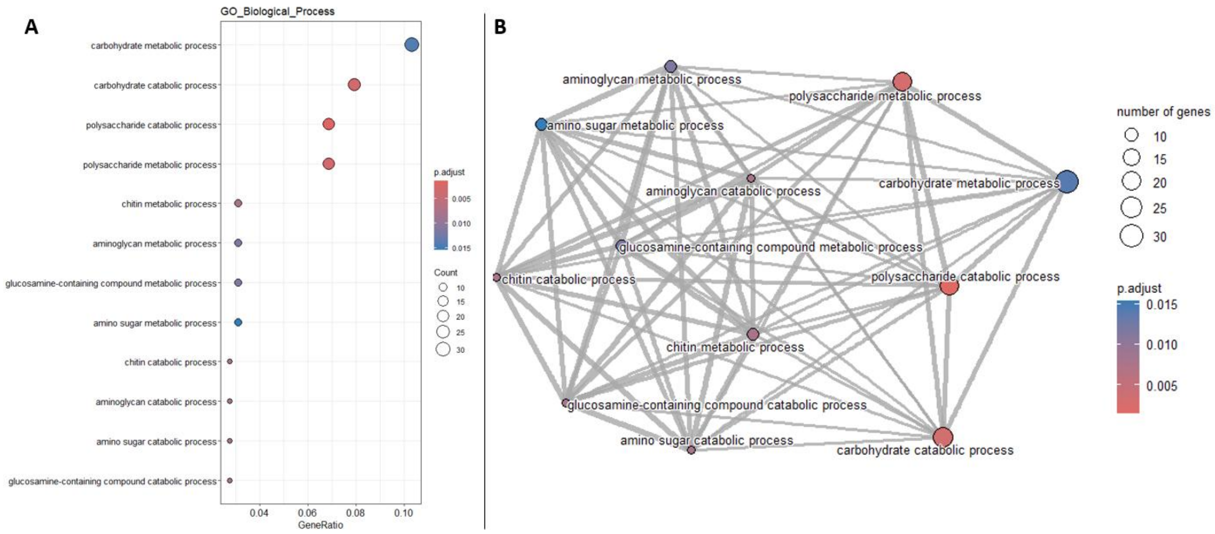


Supplementary Figure 1: Functional enrichment of genes overexpressed in response to *A. brassicicola* and *R. solani*.

The analysis was based on the 392 genes overexpressed in response to *Alternaria brassicicola* and *Rhizoctonia solani*, but not differentially expressed in response to *Globisporangium ultimum* or during the self-confrontation. Dot plot (A) and Enrichment Map (B) showing the 30 most significantly enriched GO (Gene Ontology) biological processes.

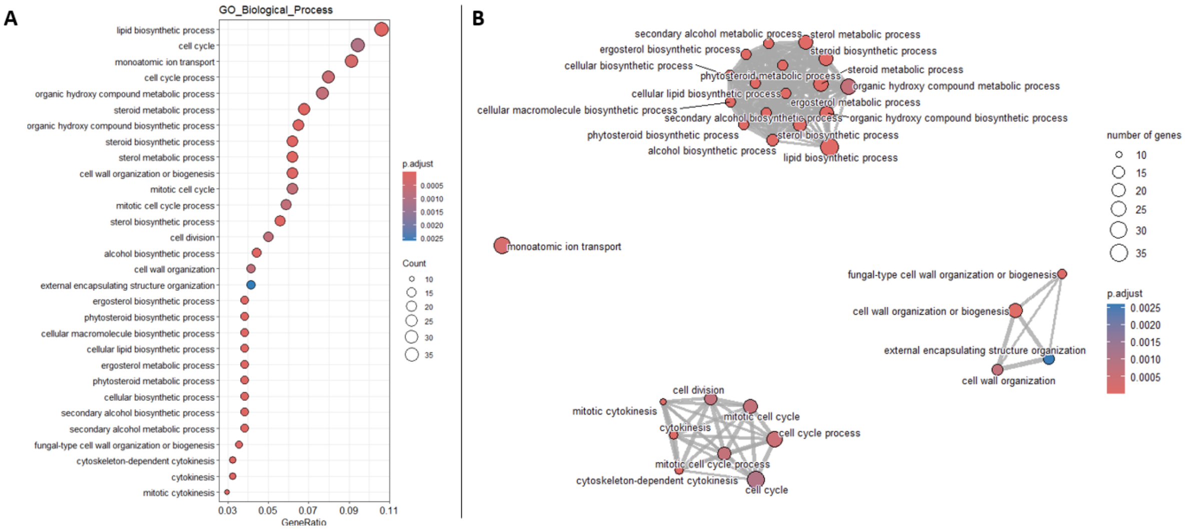


Supplementary Figure 2: Functional enrichment of genes overexpressed in response to *A. brassicicola*.

The analysis was based on the 807 genes overexpressed in response to *Alternaria brassicicola* and not differentially expressed in response to *Rhizoctonia solani*, *Globisporangium ultimum*, or during the self-confrontation. Dot plot (A) and Enrichment Map (B) showing the 30 most significantly enriched GO (Gene Ontology) biological processes.

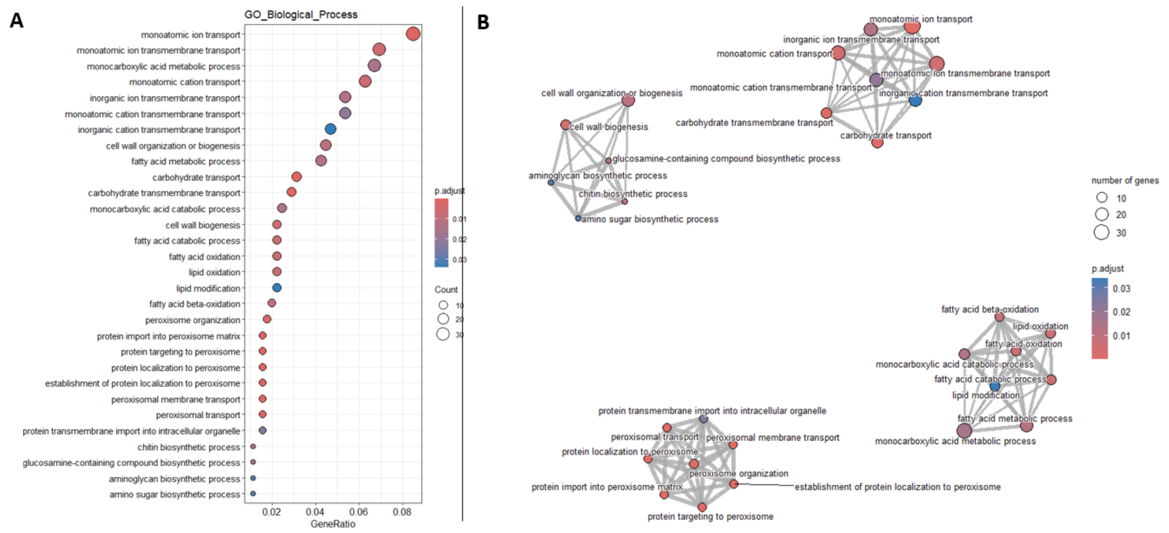


Supplementary Figure 3: Functional enrichment of genes overexpressed in response to *R. solani*. The analysis was based on the 807 genes overexpressed in response to *Rhizoctonia solani* and not differentially expressed in response to *Alternaria brassicicola*, *Globisporangium ultimum*, or during the self-confrontation. Dot plot (A) and Enrichment Map (B) showing the 30 most significantly enriched GO (Gene Ontology) biological processes.

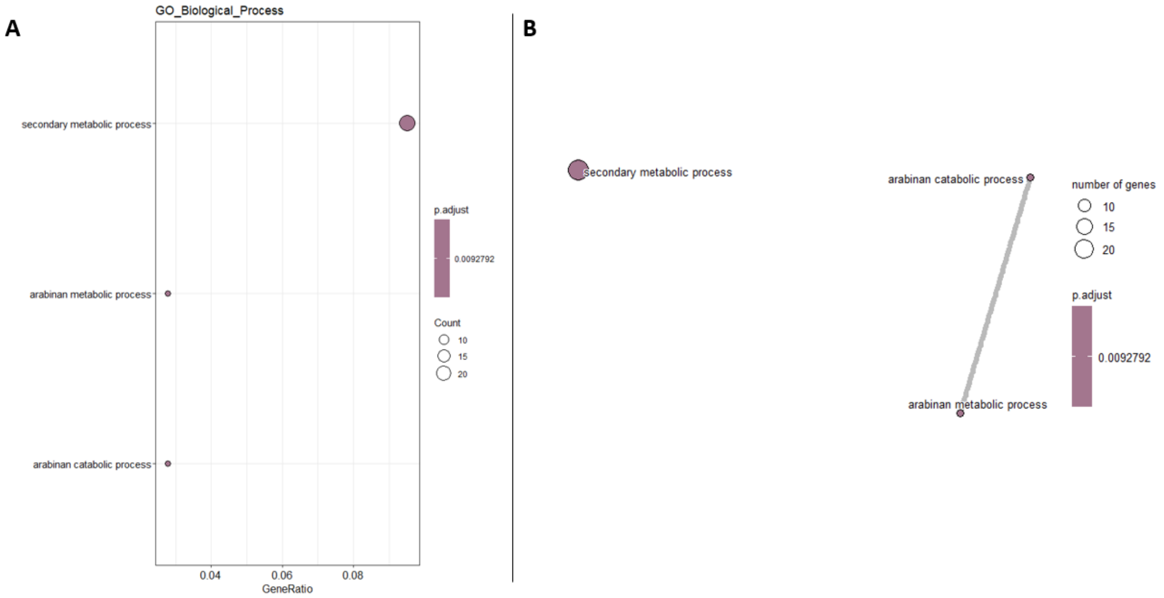


Supplementary Figure 4: Functional enrichment of genes overexpressed in response to *G. ultimum*. The analysis was based on the 807 genes overexpressed in response to *Globisporangium ultimum* and not differentially expressed in response to *Alternaria brassicicola*, *Rhizoctonia solani*, or during the self-confrontation. Dot plot (A) and Enrichment Map (B) showing the 30 most significantly enriched GO (Gene Ontology) biological processes.

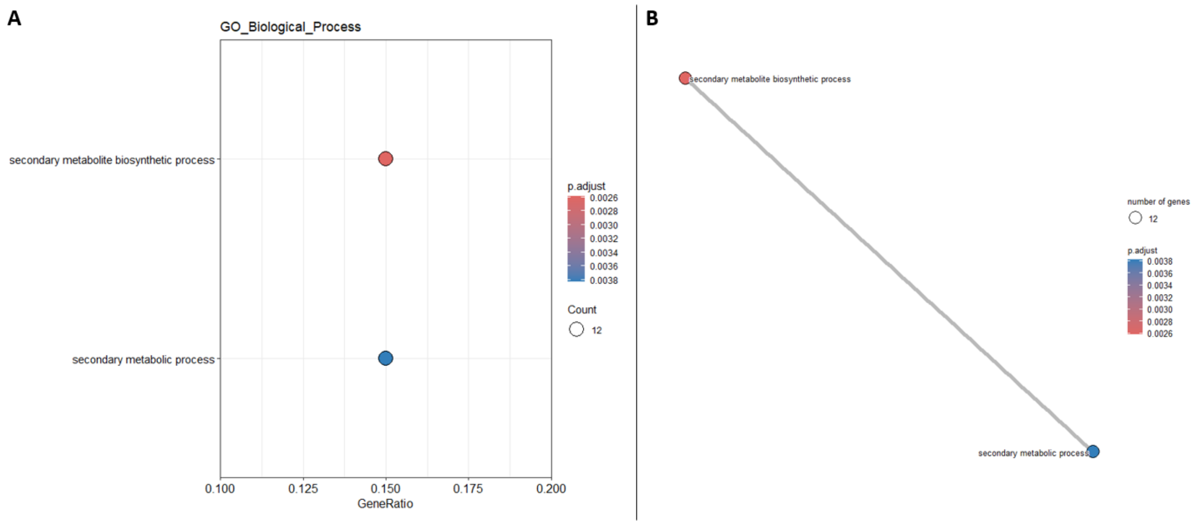


Supplementary Figure 5: Functional enrichment of genes overexpressed in response to *A. brassicicola* and *G. ultimum*.

The analysis was based on the 392 genes overexpressed in response to *Alternaria brassicicola* and *Globisporangium ultimum*, but not differentially expressed in response to *Rhizoctonia solani* or during the self-confrontation. Dot plot (A) and Enrichment Map (B) showing the 30 most significantly enriched GO (Gene Ontology) biological processes.
